# From 3D to 3D: isolation of mesenchymal stem/stromal cells into a three-dimensional human platelet lysate matrix

**DOI:** 10.1101/617738

**Authors:** Dominik Egger, Ana Catarina Oliveira, Barbara Mallinger, Hatim Hemeda, Verena Charwat, Cornelia Kasper

**Author notes:** **Correspondence**: Dr. Dominik Egger, Department of Biotechnology, University of Natural Resources and Life Science, Vienna, Austria, Muthgasse 18, 1190 Vienna, Austria.

## Abstract

Mesenchymal stem/stromal cells (MSCs) are considered an important candidate in cell therapy and tissue engineering approaches. The culture of stem cells in a 3D environment is known to better resemble the *in vivo* situation and to promote therapeutically relevant effects in isolated cells. Therefore, the aim of this study was to develop an approach for the isolation of MSCs from adipose tissue into a 3D environment. Furthermore, the use of cryoprotective medium for cryopreservation of whole adipose tissue was evaluated. For the isolation of MSCs, a novel human platelet lysate-based hydrogel was used as matrix and the migration, yield, viability and metabolic activity of cells from the 3D matrix were compared to cells from 2D explant culture. Also, the surface marker profile and differentiation capacity of MSCs from the 3D matrix were evaluated and compared to MSCs from isolation by enzymatic treatment. We found that cryopreservation of whole adipose tissue is feasible, and therefore adipose tissue can be stored and is available for MSC isolation on demand. Also, we demonstrate the isolation of MSCs into the 3D matrix and that cells from this condition display a similar phenotype and differentiation capacity like MSCs derived by traditional isolation procedure. The presented approach allows, for the first time, to isolate MSCs directly into a soft 3D hydrogel environment, avoiding any contact to a 2D plastic culture surface.

**Significance Statement:** In this paper we present a new method for the isolation of mesenchymal stem cells. Usually, these cells grow on two-dimensional plastic surfaces which is far away from their physiologic environment. Our new method allows for the first time the direct outgrowth of cells from primary tissue into a three-dimensional environment, avoiding any contact to a two-dimensional plastic surface. In future, this will allow an entirely three-dimensional in vitro cultivation of stem cells. Using 3D isolated cells will probably also increase the physiologic relevance of in vitro models.

## 1 Introduction

Mesenchymal stem/stromal cells (MSCs) are an important source for cell therapy and tissue engineering applications. They comprise a heterogeneous cell population derived from the mesenchyme, and are predominantly isolated from bone marrow^1^, adipose tissue^2^ and birth-associated tissues and fluids^3–5^. But they are also present in a variety of other tissues, such as tendon^6^, ligaments^7^ or skin^8^. They are defined by plastic adherence, trilineage differentiation (adipogenic, chondrogenic, osteogenic) and a specific surface marker expression profile (CD105^+^, CD73^+^, CD90^+^, CD14^-^, CD19^-^, CD34^-^, CD45^-^ and HLA^-^DR^-^) ^9, 10^. The regenerative potential of MSCs is not limited to their high *in vitro* proliferation potential and their ability to differentiate into adipocytes, chondrocytes and osteoblasts. Also, *in vitro* differentiation to neurons^11^, cardiomyocytes^12^ and corneal epithelial cells^13^ was observed. Effects related to injury repair were also reported, such as migration to injury sites^14^, immunomodulatory and anti-inflammatory properties mediated by cellular crosstalk, secretion of trophic factors^15^, angiogenesis^16^ and anti-scarring effects^17^.

Adipose tissue represents an easily accessible and ethically not questionable source of MSCs. The cells are usually isolated by mechanical dissociation and subsequent enzymatic digestion of the tissue with collagenase^18^. However, it was shown that commercially available collagenase products may contain endotoxins^19, 20^ and other impurities, including unwanted proteases, since they are rarely purified products. Also, cells that undergo enzymatic digestion were demonstrated to show decreased viability due to lytic activity of the enzyme^21^. To avoid damage by enzymatic digestions or impurities, MSCs were also isolated by explant culture of adipose tissue^22, 23^. Nevertheless, the final step in both isolation procedures is the selection of MSCs by adherence to a 2D plastic surface. However, the isolation procedure and the culture conditions during isolation can select subpopulations of MSCs and affect their function and potency^24^ Furthermore, cultivating MSCs in a 3D environment, either on scaffolds or scaffold-free as aggregates, is known to better reflect the physiological environment of MSCs and to have effects on cellular behavior and functionality^25–27^ Still, to the best of our knowledge, only one procedure for the isolation of MSCs into a 3D environment, avoiding selection by plastic adherence on a 2D surface, is available^28^. Papadimitropoulos et al. developed this procedure to isolate cells from a bone marrow aspirate on a perfused 3D scaffold to avoid 2D plastic adherence. Although this method allows for a streamlined MSC expansion in a bone marrow-like environment, it is limited to hard and porous scaffolds, such as ceramics. Therefore, in this study we present an approach for the direct isolation of MSCs from adipose tissue into a soft 3D environment. For this, we used a hydrogel matrix prepared from polymerized human platelet lysate (hPL). This hydrogel has the advantage that it serves, both as adhesion matrix and nutrient supply and it was already been shown to be suitable for 3D cell culture of MSCs^29, 30^. Furthermore, it is of increasing interest to have human adipose tissue for the isolation of MSCs available at any time. Since donor tissue is not always available on demand, strategies for the long term storage of human adipose tissue are required. The cryopreservation of liposuction aspirates from adipose tissue and isolation of MSCs by enzymatic treatment from long term cryopreserved adipose tissue was demonstrated before^31, 32^ However, we present an approach for the cryopreservation of whole adipose tissue portions for isolation of MSCs by explant cultivation.

## 2 Materials and methods

### 2.1 Freezing and thawing of adipose tissue

The adipose tissue used in this study was derived from abdominal plastic surgery of four different donors (female, 28 – 58 y.o.), as described before^33^. The tissue was manually cut into pieces of approximately 125 mm^3^, as a maximum side length of 5×5×5 mm was intended. Afterwards, 1 g (corresponding to approximately 8 pieces) of tissue was transferred to a vial for cryopreservation (Greiner Bio-One, Kremsmünster, Austria), covered with medium for cryopreservation composed of MEM alpha (Thermo Fisher Scientific, Waltham, MA, USA) supplemented with 12.5 % hPL (PLSolution; PL BioScience, Aachen, Germany) and 10 % dimethyl sulfoxide (DMSO; Sigma Aldrich, St. Louis, MO, USA) or left without medium and subsequently cooled down at 1 °C/min to −80 °C. After 24 h, the tissue was stored in liquid nitrogen (−196 °C). Upon thawing, the vials were placed in a 37 °C water bath for 2 min, the medium was removed and the pieces either transferred to a standard cell culture plate or embedded into a gel comprised of polymerized hPL as described in section 2.2 and 2.3.

### 2.2 Isolation via 2D explant culture

For the isolation via 2D explant culture, one thawed tissue piece (cryopreserved without medium) was transferred to tissue culture treated 24-well plates, and incubated for 1 h at 37 °C to let the tissue attach to the surface. Some experiments were performed in a 6-well plate. In this case, 3 pieces of tissue were transferred to one well. Afterwards, the tissue was covered with expansion medium composed of MEM alpha with 10 % hPL, 0.5 % gentamycin (Lonza, Basel, Switzerland) and 1 U/ml PLSupplement (heparin; PL BioScience GmbH) and incubated in a standard incubator. For better comparison, the tissue was covered with the same volume of medium as the volume of PLMatrix (PL BioScience GmbH, Aachen, Germany) used during 3D explant culture (refer to section 2.3), and no medium changes were performed. After 14 days, the cells were detached by accutase (Sigma Aldrich) treatment (15 min incubation at 37 °C), and the total cell number was determined by trypan blue staining and manual counting with a hemocytometer. The standard enzymatic isolation procedure from adipose tissue was performed as described before^33^.

### 2.3 Isolation via 3D explant culture

For the isolation via 3D explant culture, adipose tissue was embedded between two layers of a hPL-based gel, which was composed of 10 % reconstituted lyophilized PLMatrix (PL BioScience GmbH, Aachen, Germany) in MEM alpha with 0.5 % gentamycin. First, a bottom layer was added to 6- or 24-well plates and polymerized for 1 h at 37 °C in a standard incubator. As for the 2D explant culture, either 3 or 1 thawed tissue pieces (cryopreserved without medium) were transferred on top of the bottom layer in a 6- or 24-well plate, respectively, covered with a top layer of PLMatrix and incubated at 37 °C for up to 14 days.

### 2.4 Harvesting of MSCs from PLMatrix

The cells that were cultured in the PLMatrix were harvested by combining mechanical dissociation and enzymatic digestion of the extracellular matrix (ECM). The gel was dissociated by suction through the needle of a 5 ml syringe (both Braun, Kronberg im Taunus, Germany), and the liquid gel, together with the tissue, was transferred to a centrifugation tube. The well was rinsed with phosphate-buffered saline (PBS; Sigma Aldrich), transferred to the same tube, and the tube was centrifuged at 500 g for 5 min. After removal of the supernatant and the adipose tissue, the cell pellet was resuspended in 2 ml of 2 mg/ml collagenase IA (Sigma Aldrich) in PBS and incubated for 1 h at 37 °C on a horizontal shaker at 100 rpm. Afterwards, the tube was centrifuged at 500 g for 5 min and the pellet resuspended in expansion medium. The total cell number was then determined by trypan blue staining and manual cell counting with a hemocytometer.

### 2.5 Differentiation

To determine the adipogenic and osteogenic differentiation capacity, 4,000 cells per cm^2^ in passage 2 were seeded on a 12 well plate coated with fibronectin (2 μg/cm^2^; Sigma Aldrich) and allowed to grow confluent. For chondrogenic differentiation, 2.5 x 10^5^ cells were seeded into a 15 ml tube and centrifuged (300 xg, 5 min) to form an aggregate. Then, in all conditions, medium was changed to adipogenic, chondrogenic (NH AdipoDiff Medium or NH ChondroDiff Medium, both Miltenyi Biotech, Bergisch Gladbach, Germany, with 0.5 % gentamycin) or osteogenic differentiation medium (MEM alpha, 2.5 % HPL, 0.5 % gentamycin, 1 U/ml heparin, 5 mM beta-glycerolphosphate, 0.1 μM dexamethasone and 0.2 mM L-ascorbate-2-phosphate, all Sigma Aldrich), respectively. Cells were cultivated for 21 days, and the medium was changed every 2 – 3 days.

### 2.6 Histological stainings

MSCs cultivated in adipogenic medium were stained for lipid vacuoles with Oil Red O, after fixation with 4 % paraformaldehyde (PFA, both Sigma). To confirm chondrogenic differentiation, the aggregates were fixed with 4 % PFA, embedded in paraffin and stained for glycosaminoglycans with Alcian Blue (Sigma) according to routine histology protocols. MSCs cultivated in osteogenic medium were fixed with 96 % ethanol and stained for calcium with Alizarin Red (Sigma). Cells on day 0 and day 21 cultivated in expansion medium served as control. All stainings were performed as described before in detail^27^

### 2.7 Preparation of histological sections

To prepare histological sections of PLMatrix, the matrix with adipose tissue and migrated cells was fixed with 4 % PFA for 24 h, embedded in paraffin using a Shandon Tissue Excelsior (Thermo Fisher Scientific) and cut with a rotating microtome (Thermo Fisher Scientific). Subsequently, the sections were stained with hematoxylin (Richard Allan Scientific) and eosin (Carl Roth) in ddH_2_O and dehydrated, before sections were covered with DPX mounting medium (Sigma Aldrich).

### 2.8 Nuclear staining in PLMatrix

Prior to staining the cell nuclei with 4’,6-diamidin-2-phenylindol (DAPI), the samples were fixed with 4 % PFA (both Sigma Aldrich). The cells from standard cell culture plastic surfaces were rinsed with PBS, whereas the PLMatrix samples were not rinsed. Afterwards, the samples were covered with DAPI 1 μl/ml in PBS with 0.1 % Triton X (Sigma Aldrich) and incubated for 30 min at room temperature. Subsequently, the staining solution was discarded, PBS added and the staining documented by fluorescence microscopy (Leica, DMIL LED, EL6000).

### 2.9 Live/dead staining

Viability of cells was visualized with calcein-acetoxymethyl ester (AM) and propidium iodide (PI) staining. Briefly, the samples were covered with Calcein-AM (2 μM) and PI (8 μM) in PBS and incubated for 30 min at 37 °C. The cells from standard cell culture plastic surfaces were rinsed with PBS prior to incubation. After incubation. all samples were rinsed with PBS and the staining documented by fluorescence microscopy.

### 2.10 Metabolic activity

To assess the metabolic activity of cells after isolation, the cells were first harvested and incubated with collagenase to obtain single cells as described in section 2.4. Afterwards, cells were seeded at 1 x 10^4^ cells/cm^2^ on standard cell culture plates in expansion medium and incubated overnight at 37 °C. Subsequently, a resazurin-based *in vitro* toxicology assay kit (TOX8) was applied (Sigma Aldrich) and fluorescence intensity was measured with a plate reader (Tecan, Männedorf, Switzerland) according to the manufacturer’s instructions. Expansion medium served as a control.

### 2.11 Phenotyping

To determine MSC surface marker expression, the cells of one donor were detached by accutase treatment and stained with a human MSC phenotyping kit and an anti-HLA-DR antibody (both Miltenyi Biotec) according to the manufacturer’s instructions. In this kit, the antibodies for the negative markers (CD14, CD20, CD34 and CD45) are labeled with the same fluorophore to generate a negative marker panel. According to the manufacturer’s manual, an approximately 10-fold increase of the fluorescence intensity of the negative markers is expected for negative samples, as compared to the isotype control. The stained cells were resuspended in a suitable volume of flow cytometry buffer (0.5 % fetal bovine serum, 2 mM EDTA in PBS), and acquisition was carried out on a BD FACS Canto II (Franklin Lakes, New Jersey, US). Between 1–5 x 10^4^ gated events were recorded per sample. Subsequent analysis was performed with Kaluza Flow Cytometry software (version 1.3, Beckman Coulter, Brea, CA, USA).

### 2.12 Statistical analysis

All data is expressed as mean values ± standard deviation (SD). The data was analyzed using Microsoft Excel and GraphPad Prism 6.01. Comparisons were carried out using unpaired t-tests. Values of p < 0.05 with a confidence interval of 95 % were defined as statistically significant.

## 3 Results and Discussion

### 3.1 Cryopreservation of adipose tissue

We tested if our standard medium for cryopreservation of MSCs (12.5 % hPL, 10 % DMSO in MEM alpha) would also be suitable for the cryopreservation of small pieces of human adipose tissue (approximately 125 mm^3^), and tested this condition against cryopreservation without medium or additional cryo-protectives. After thawing, the tissue was embedded in PLMatrix for the isolation of cells. The outgrowth of cells was observed in both conditions, and no significant difference was observed during 14 days (Figure 1B). The number of viable cells harvested from the PLMatrix was significantly higher after cryopreservation with medium, compared to cryopreservation without medium (1.9 ± 0.3 vs. 1.1 ± 0.1 x 10^5^ cells). The overall viability was similar after cryopreservation with and without medium (91 ± 5 % vs. 95 ± 13 %). However, the metabolic activity of cells after isolation and harvest was significantly higher (1.5 ± 0.2 fold) in cells cryopreserved without medium, compared to cells cryopreserved with medium (Figure 1E). DMSO is usually used at 5 or 10 % as a cryoprotective agent in cryomedia. Although it is known to cause damage to cells^31^, the use of 10 % DMSO (compared to 5 %) increased the recovery and viability of cells after cryopreservation^34^, which is why 10 % DMSO was used for cryopreservation in this study. However, the decreased metabolic activity might be due to remnants of DMSO in the adipose tissue. This data suggests that cryopreservation of small pieces (approximately 125 mm^3^) of whole adipose tissue with and without a cryoprotective medium is feasible. Although the cell number after thawing was higher when a cryoprotective medium was used, the omission of a cryoprotective medium containing DMSO seems to cause less cell stress and might be more cost- and time-effective in handling. However, we decided to freeze and thaw adipose tissue without cryoprotective medium for the comparison of 3D vs. 2D explant isolation in this study.

**Figure 1.**
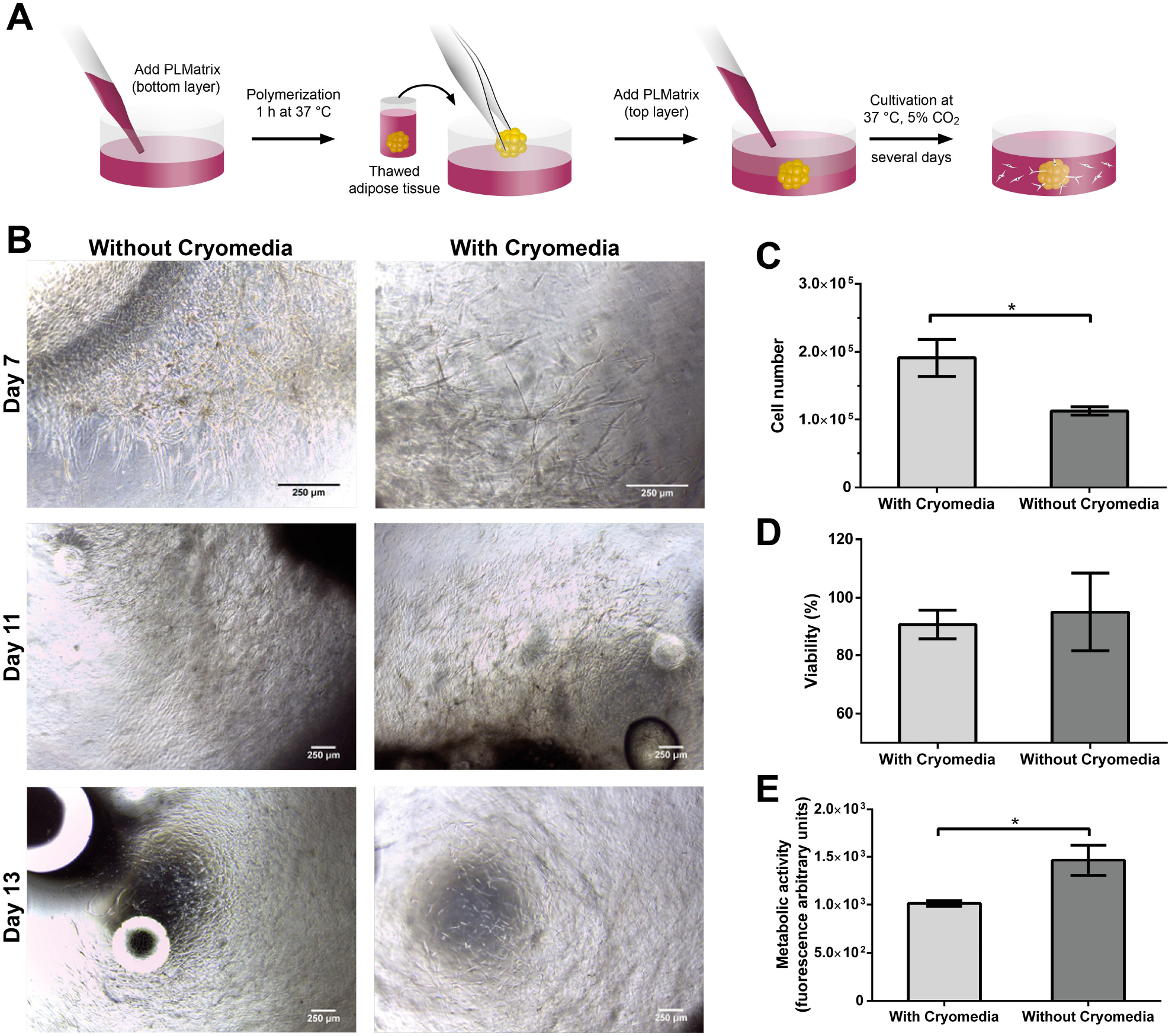
Migration and evaluation of cells from adipose tissue after cryopreservation of whole adipose tissue with and without cryoprotective medium. **(A)** Scheme of embedding adipose tissue into PLMatrix for 3D explant isolation of MSCs. **(B)** Micrographs showing the outgrowth of MSCs from adipose tissue into PLMatrix after thawing of adipose tissue without and with cryomedia at different time points. **(C)** Cell number, **(D)** viability and **(E)** metabolic activity assessed with resazurin-based TOX-8 assay of MSCs after outgrowth from adipose tissue cryopreserved without and with cryomedia into PLMatrix after 14 days of outgrowth. Data represented as average ± SD from n=3 experiments, asterisk indicate statistical significance.

### 3.2 Migration of MSCs into PLMatrix

The PLMatrix hydrogel was demonstrated to be suitable for cell culture of MSCs in prior studies^29, 30^. Since the matrix is based on polymerized hPL, the hydrogel serves not only as a structure for migration of the cells but at the same time as nutrient supply. In this study, we compared the migration of cells from adipose tissue into the PLMatrix with the migration of cells on a standard 2D surface (2D explant) over a period of 14 days. Individual cells were visible in the matrix as early as 72 h after embedding the tissue into PLMatrix (Supplementary figure 1 and supplementary video). After that, the cells continued to migrate out and proliferated until day 14, as demonstrated by histological sections of the gel and micrographs (Figure 2). As the number of harvested cells did not change significantly after 11 days, observations were discontinued at day 14. Generally, more cells could be harvested from the PLMatrix compared with 2D explant (Figure 2C). This is facilitated by the possibility to migrate from the tissue and migrate not only in two, but three dimensions. Most of the cells were found to be viable in both conditions, as confirmed by Calcein-AM and PI staining. It has been reported that the isolation of adipose-derived MSCs by explant culture increases the cell yield compared to isolation by enzymatic digestion, while the cells displayed similar comparable immunophenotypic and functional properties^22^ We found that the use of PLMatrix for the isolation of MSCs by explant culture can further increase the yield since cells can grow in all three dimensions, while occupying the same surface area. Interestingly, the histological sections displayed two distinct states of infiltrated areas (Supplementary figure 2). The more distant areas, which were infiltrated by only few cells, barely displayed collagen, as indicated by eosin staining. In comparison, areas with a higher cell density displayed higher amounts of collagen. During the harvest procedure, the collagen was not totally degraded by collagenase treatment, and single cells were found to still be attached to the collagen fibers after digestion. Therefore, optimization of the harvest procedure might further enhance the efficiency and reproducibility of isolation by 3D explant culture. Also, future approaches need to concentrate on more effective and safe alternatives to avoid the use of collagenase.

**Figure 2.**
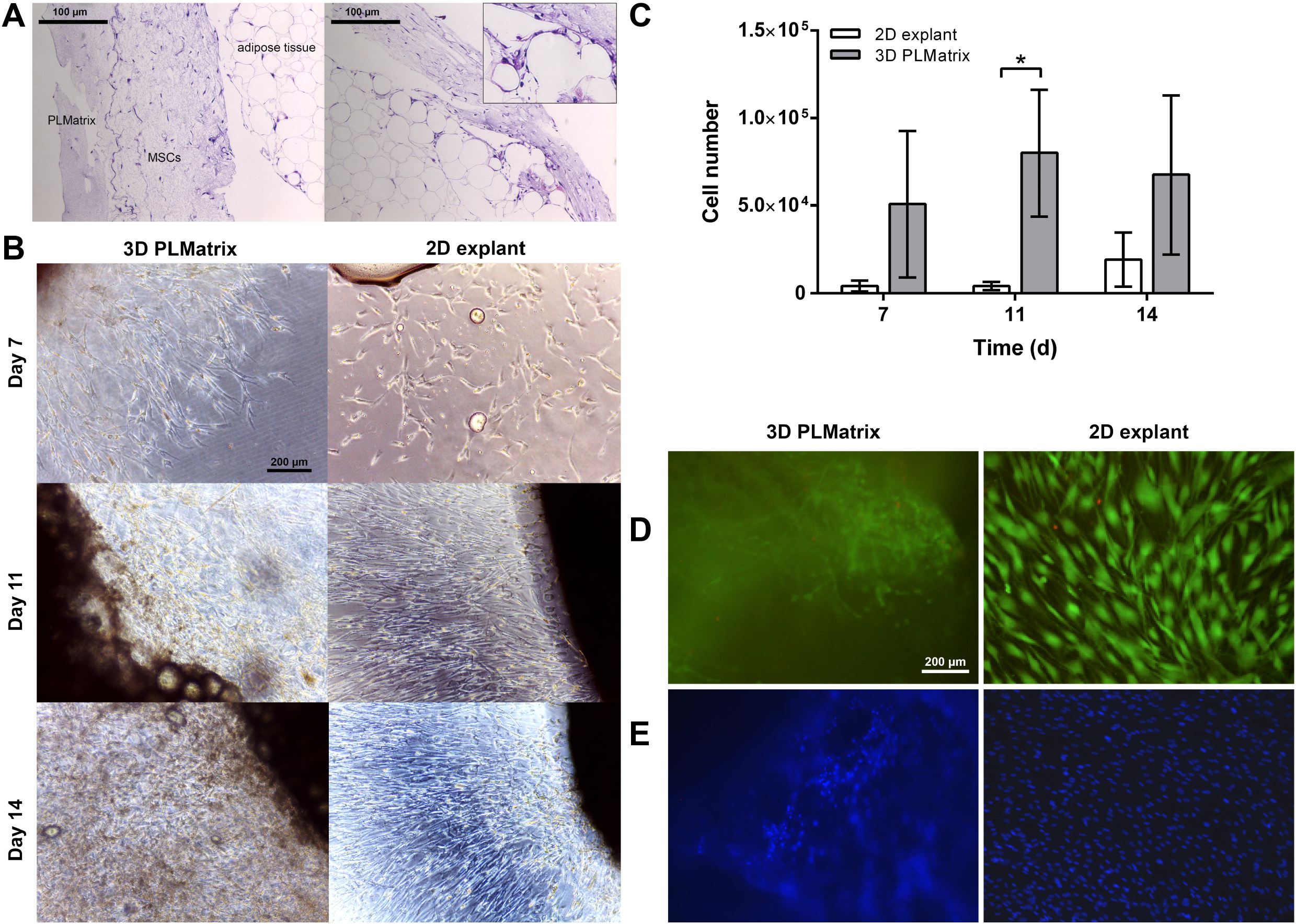
Comparison of 2D and 3D conditions for the isolation of MSCs from whole adipose tissue. **(A)** Histological sections stained with hematoxylin (blue, stains for nuclei) and eosin (pink, stains for collagen) of adipose tissue embedded in 3D PLMatrix hydrogel with MSCs migrating into the hydrogel after 11 days. **(B)** Outgrowth of MSCs from adipose tissue into 3D PLMatrix or on a 2D plastic surface (2D explant) after 7, 11 and 14 days of culture. **(C)** Amount of MSCs harvested from PLMatrix or 2D plastic surface after 7, 11 and 14 days of outgrowth from adipose tissue. **(D)** Calcein-AM and PI and **(E)** DAPI stain of MSCs in PLMatrix and on 2D plastic surface on day 14 after outgrowth from adipose tissue. Data represented as average ± SD from n=3 experiments, asterisk indicate statistical significance.

Another important factor for future optimization is matrix stiffness, which impacts cell migration, differentiation and ECM production in a 3D environment^35, 36^. The Young’s modulus of PLMatrix was determined prior to use in cell culture and was found to be 0.1 kPa (data not shown), which is close to clinical values for soft tissues. For example, bone marrow has a Young’s modulus of 0.3 kPa^37^, adipose tissue around 1 – 4 kPa^38, 39^ and viable human MSCs around 1 kPa^40^. Matrix composition of the surrounding tissue highly affects the elasticity, and major matrix proteins like collagen are known to increase the Young’s modulus. As described above, two distinct states of infiltrated areas were found in the histological sections (Supplementary Image 2). These observations indicate that the cells started to remodel the hydrogel over time, presumably adjusting the Young’s modulus to generate a more physiologic matrix. Some previous studies have shown that MSCs display an increased proliferation in hydrogels of higher stiffness^35, 36, 41^. Also, differentiation of MSCs is known to be regulated by substrate stiffness^42^ For example, a stiff substrate was shown to induce osteogenic differentiation, whereas a soft substrate induced neuronal differentiation^36^. Therefore, future work will focus on the development of a tunable hPL-based hydrogel with an increased Young’s modulus.

### 3.3 Characterization of MSCs derived by 3D explant isolation

To characterize the cells derived from 3D explant isolation, the differentiation capacity and phenotypic properties of the cells were determined. For this, the cells from 3D explant isolation were cultivated in adipogenic, chondrogenic and osteogenic differentiation media. Cells isolated in PLMatrix were found to differentiate into all three lineages, as confirmed by histological stainings (Figure 3, supplementary figures 3 to 5). Furthermore, they displayed the same differentiation potential as cells derived from the traditional 2D enzymatic isolation procedure. To characterize the surface marker expression, cells harvested from standard enzymatic isolation procedure (2D control), 2D explant procedure and PLMatrix were stained for CD105, CD73, CD90, CD14, CD19, CD34, CD45 and HLA-DR and analyzed by flow cytometry. The isolated cells were found to be positive for CD73, CD90 and CD105 and negative for CD14, CD20, CD34, CD45 and HLA-DR which is in accordance with the minimal criteria for the characterization of MSCs^9^ The surface marker profile and average marker expression, as compared to the isotype control, were similar across all conditions (Figure 3). Only the expression of CD73 was found to be altered in cells from 2D explant and PLMatrix, as compared to the 2D control. CD73 is expressed on a variety of cell types and known to be involved in many processes such as the adaption to hypoxia or response to inflammation^43^. Furthermore, a recent publication discussed CD73 as a universal marker for the purification of MSCs after isolation^44^ In this study, a CD29^+^CD54^+^CD31^-^CD45^-^ population isolated from rat bone marrow was identified as MSCs with high colony-forming efficiency and trilineage differentiation capacity. These cells were identical to a population isolated by positive selection of CD73 and uniformly expressed other characteristic MSC markers (CD29, CD44, and CD90). In conclusion, the cells isolated in PLMatrix were found to differentiate into adipogenic, chondrogenic and osteogenic lineage, and to express a MSC-like surface marker profile. Furthermore, differentiation potential and surface marker profiler was similar to MSCs from the traditional 2D enzymatic isolation procedure. Consequently, these cells were considered as MSCs.

**Figure 3.**
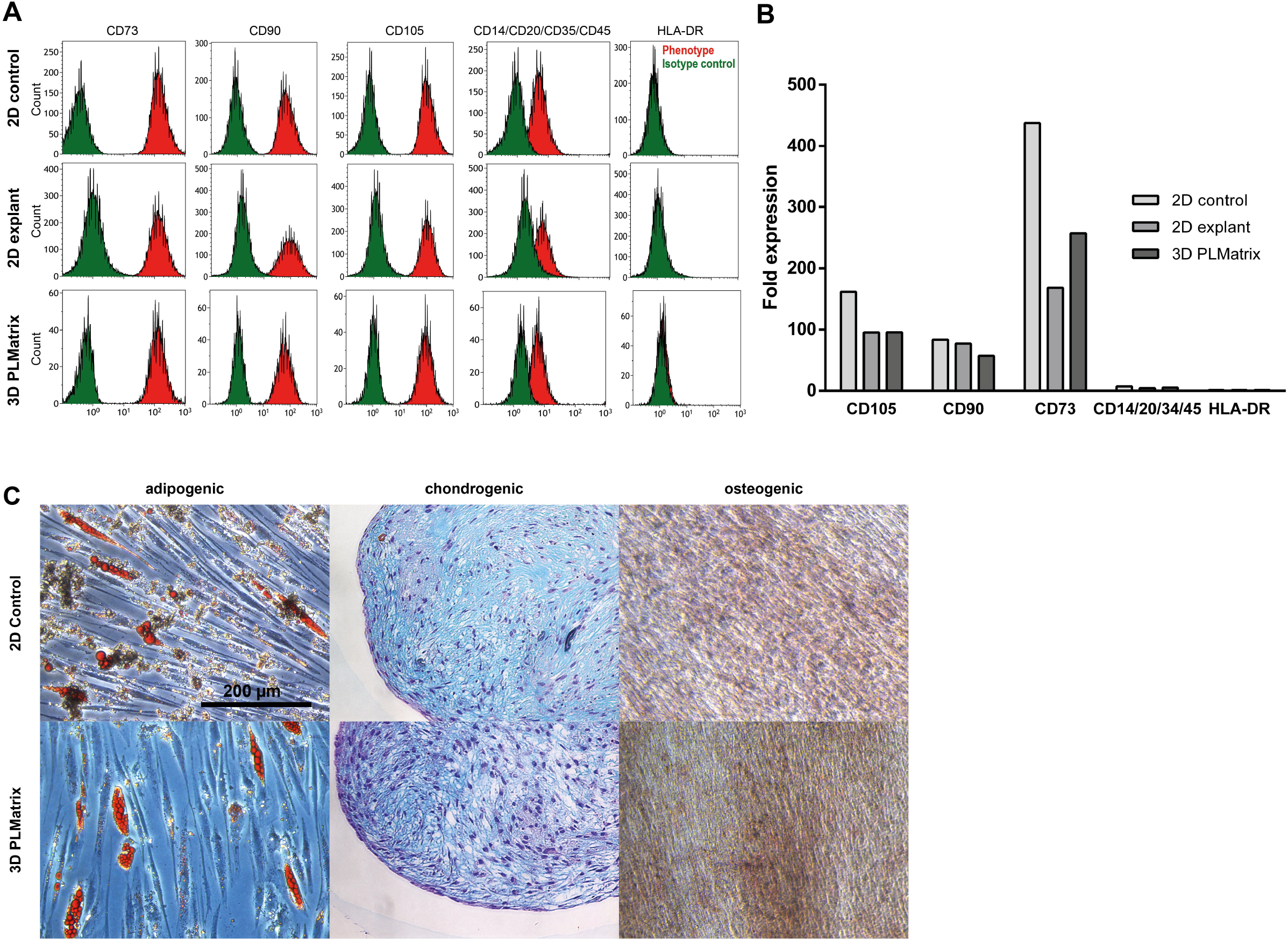
Characterization of cells that migrated from adipose tissue into PLMatrix. **(A)** Surface marker analysis of MSCs from standard enzymatic isolation procedure (2D control), isolation from tissue explants on a 2D plastic surface (2D explant) and isolation from tissue explants in the 3D PLMatrix. **(B)** Average x-fold expression of surface markers of phenotype normalized to isotype control. **(C)** Micrographs of cells cultivated in adipogenic, chondrogenic or osteogenic differentiation medium for 21 days, stained with Oil Red O, Alcian Blue and Alizarin Red, respectively. Data of flow cytometric analysis represents ≥ 10.000 gated events.

**Table 1:** Surface marker analysis of MSCs from standard enzymatic isolation procedure (2D control), isolation from tissue explants on a 2D plastic surface (2D explant) and isolation from tissue explants in the 3D PLMatrix. Percentage of positive stained events compared to the isotype control. Data represents ≥ 10.000 gated events.

| Surface Marker | 3D PLMatrix | 2D explant | 2D control |
| --- | --- | --- | --- |
| CD73 | 99.8 | 99.7 | 99.9 |
| CD90 | 99.8 | 99.7 | 99.9 |
| CD105 | 99.7 | 99.5 | 99.9 |
| CD14/CD20/CD35/CD45 | 47.0 | 52.3 | 34.9 |
| HLA-DR | 0.2 | 0.2 | 0.1 |

## 4 Concluding remarks

In this study, we found that small pieces of whole adipose tissue can be frozen and thawed either with a standard cryoprotective medium or even without medium. The use of medium with DMSO as a cryoprotective compound increased the yield, but decreased the metabolic activity of cells. Since adipose tissue is often not available on demand, this procedure can potentially be used for long term storage, however, the quality of MSC after longer storage time will have to be tested in the future. Moreover, to the best of our knowledge, we demonstrate the first approach for the direct isolation of MSCs from tissue into a soft 3D environment, avoiding any contact to a 2D surface. This process might enable to cultivate MSCs in a physiologic 3D environment for the entire period of *in vitro* culture. With the use of PLMatrix hydrogel the 3D environment serves as adhesion matrix and nutrient supply, both of which are promoting cell migration into the matrix. The development of a tunable hPL-based hydrogel with adjustable Young’s modulus is planned, in order to increase proliferation during 3D explant isolation, and thus the cellular yield. The cells harvested from the matrix were characterized towards the surface marker profile and differentiation capacity, and were found to fulfill the minimal criteria for MSCs. Traditional isolation methods based on enzymatic treatment and selection by 2D plastic surfaces do not represent a physiologic environment. Since 3D culture often leads to a behavior that is more representative of the *in vivo* situation, the isolation into a 3D environment avoiding any contact with 2D plastic surfaces might contribute to a more physiologic behavior of MSCs. However, the therapeutic potential of MSCs derived by 3D explant isolation remains to be evaluated by functional assays.

## 7 Data Availability Statement

The data that support the findings of this study are available from the corresponding author upon reasonable request.

## Author contributions

Dominik Egger: conception and design, collection and/or assembly of data, data analysis and interpretation, manuscript writing.

Ana Catarina Oliveira: collection and/or assembly of data

Barbara Mallinger: collection and/or assembly of data

Hatim Hemeda: provision of study material, conception and design

Verena Charwat: collection and/or assembly of data, data analysis and interpretation

Cornelia Kasper: conception and design, final approval of manuscript

