## Supplementary figures and images for "From 3D to 3D: isolation of mesenchymal stem/stromal cells into a three-dimensional human platelet lysate matrix"

### Supplementary figure 1

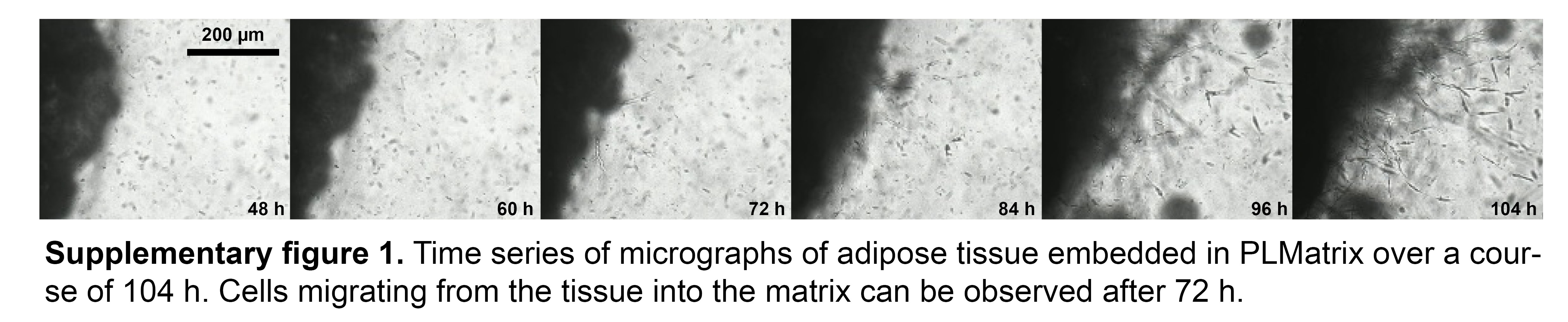

### Supplementary figure 2

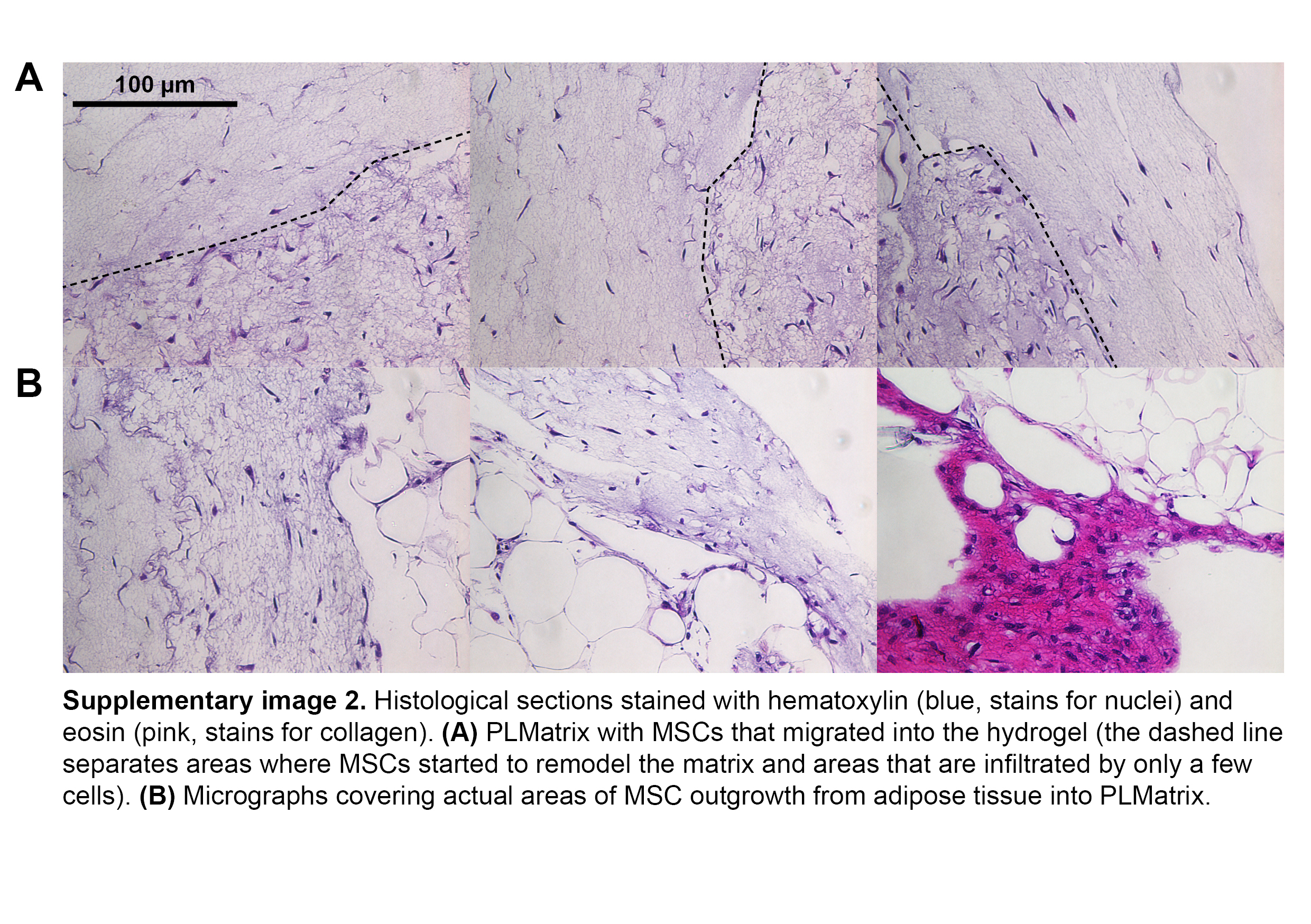

### Supplementary figure 3

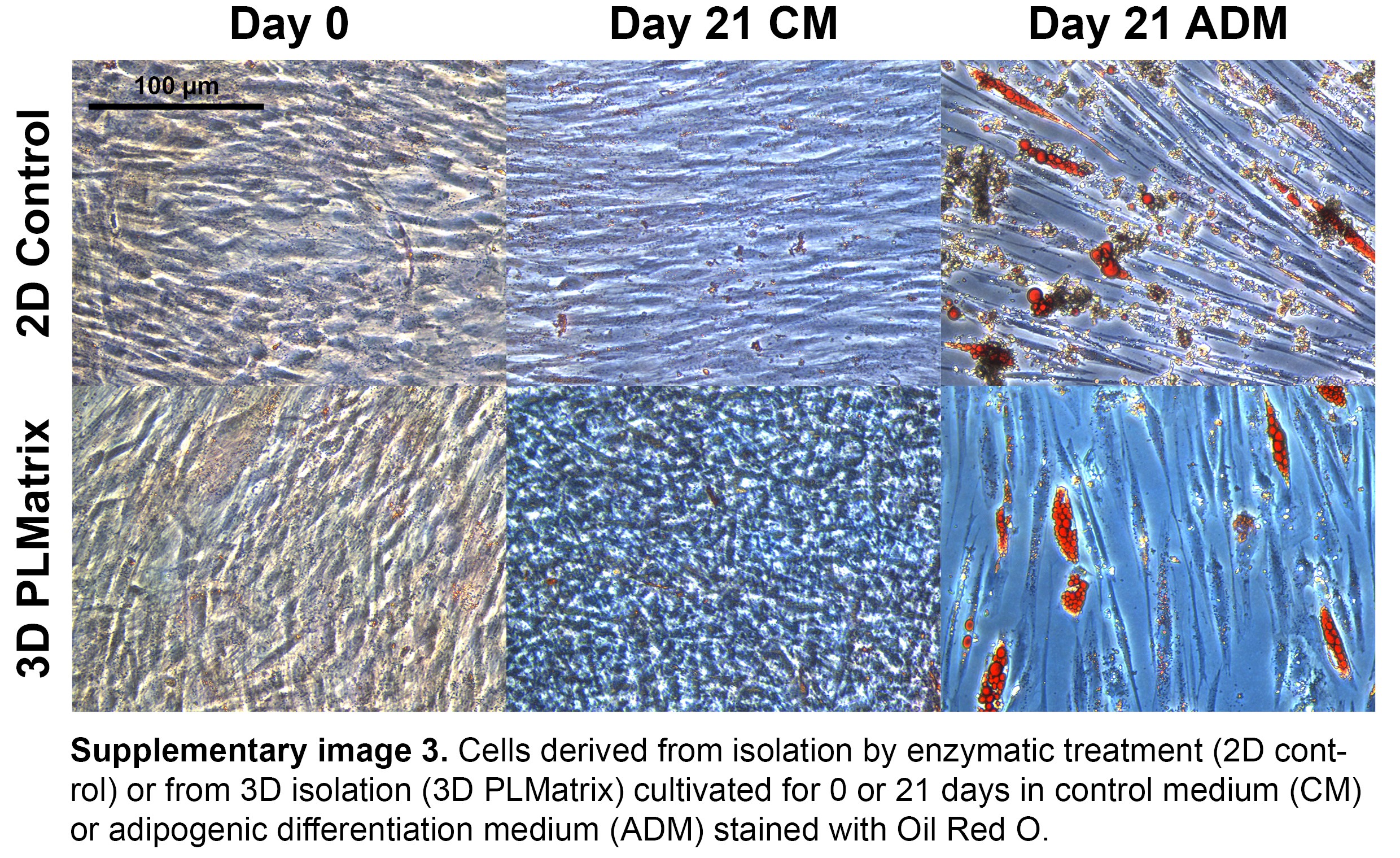

### Supplementary figure 4

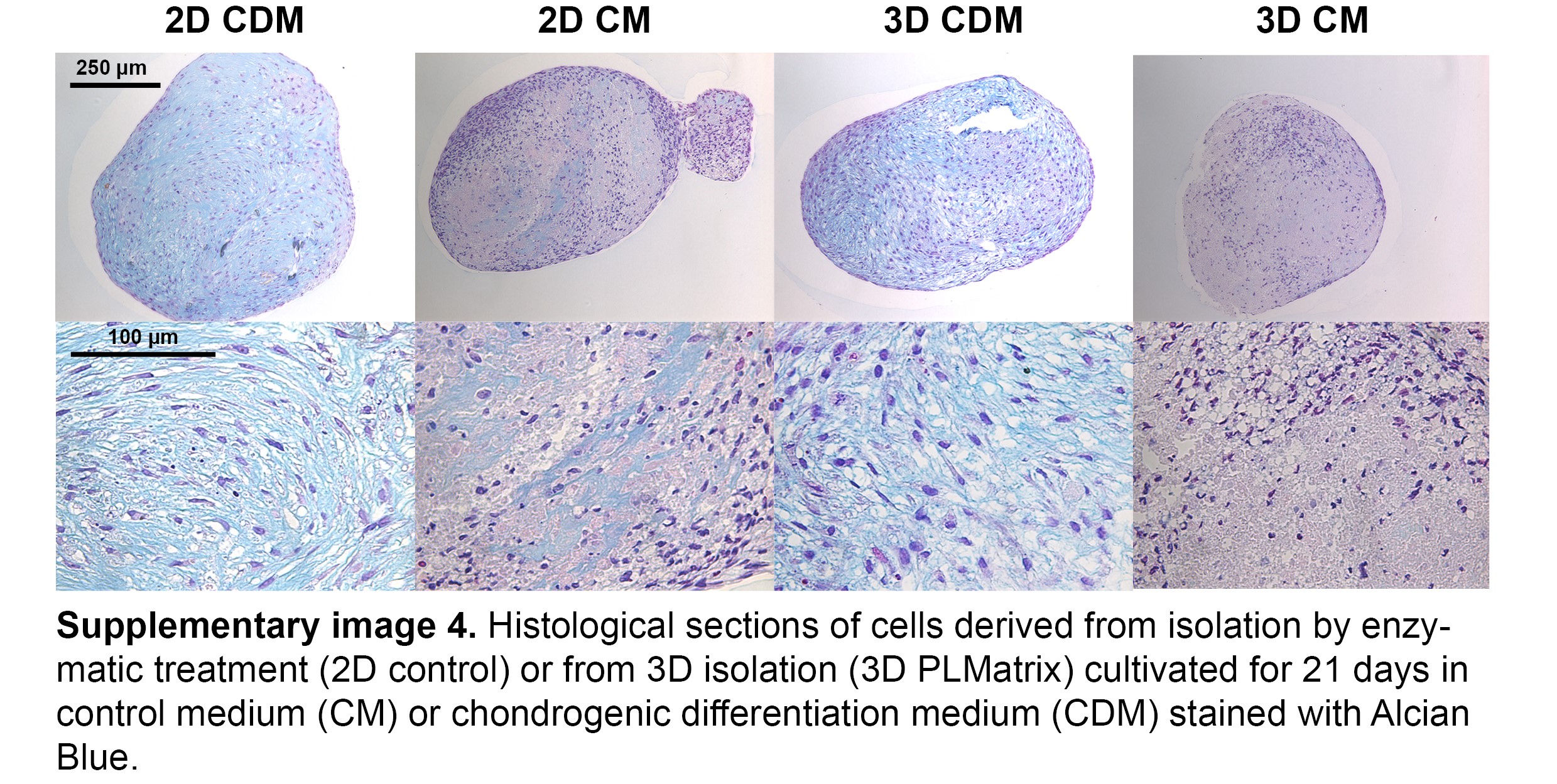

### Supplementary figure 4

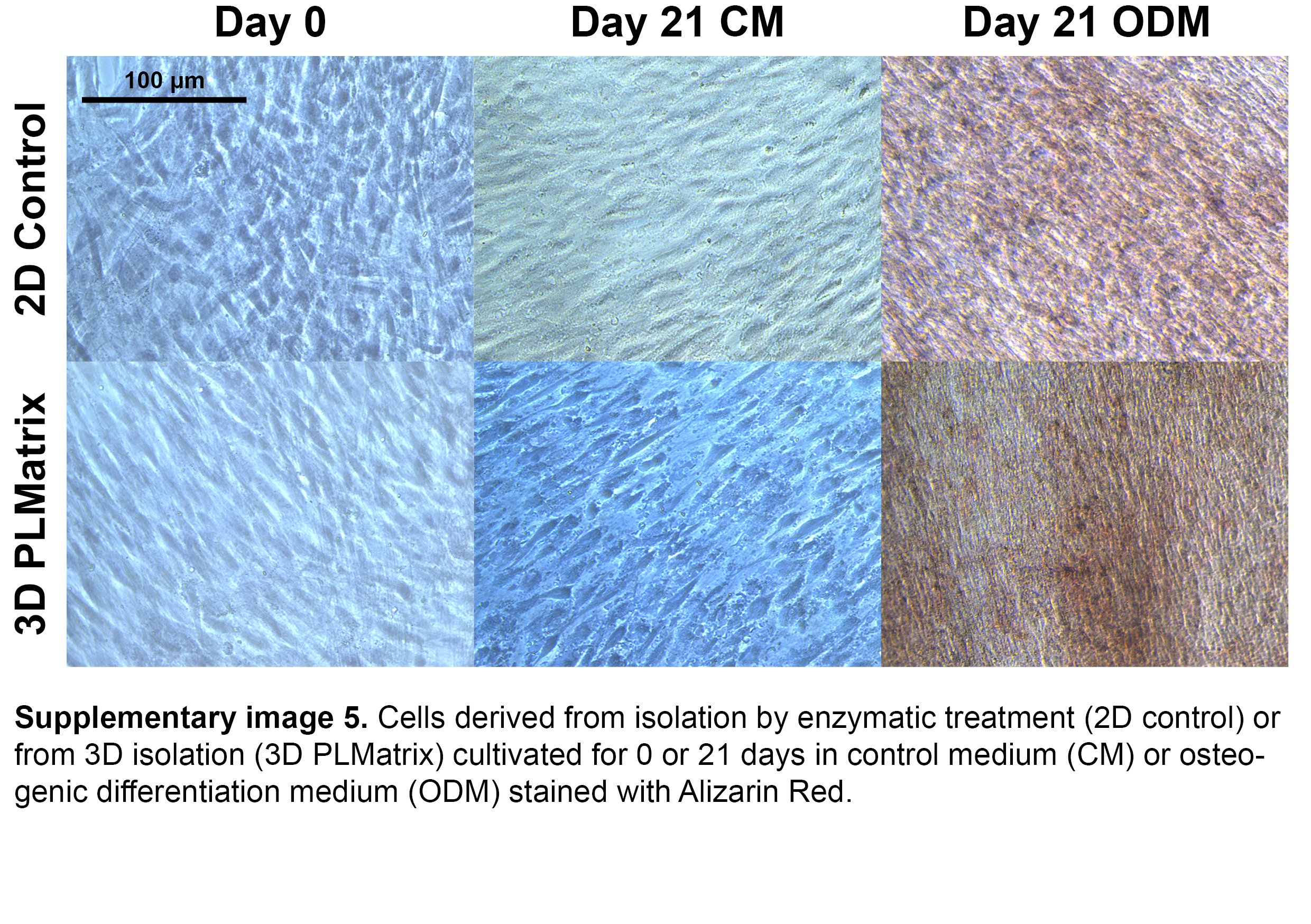
