## Supplementary figures for "From 3D to 3D: isolation of mesenchymal stem/stromal cells into a three-dimensional human platelet lysate matrix"

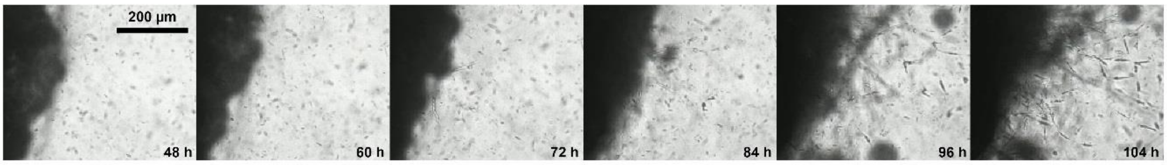


**Supplementary figure 1.** Time series of micrographs of adipose tissue embedded in PLMatrix over a course of 104 h. Cells migrating from the tissue into the matrix can be observed after 72 h.

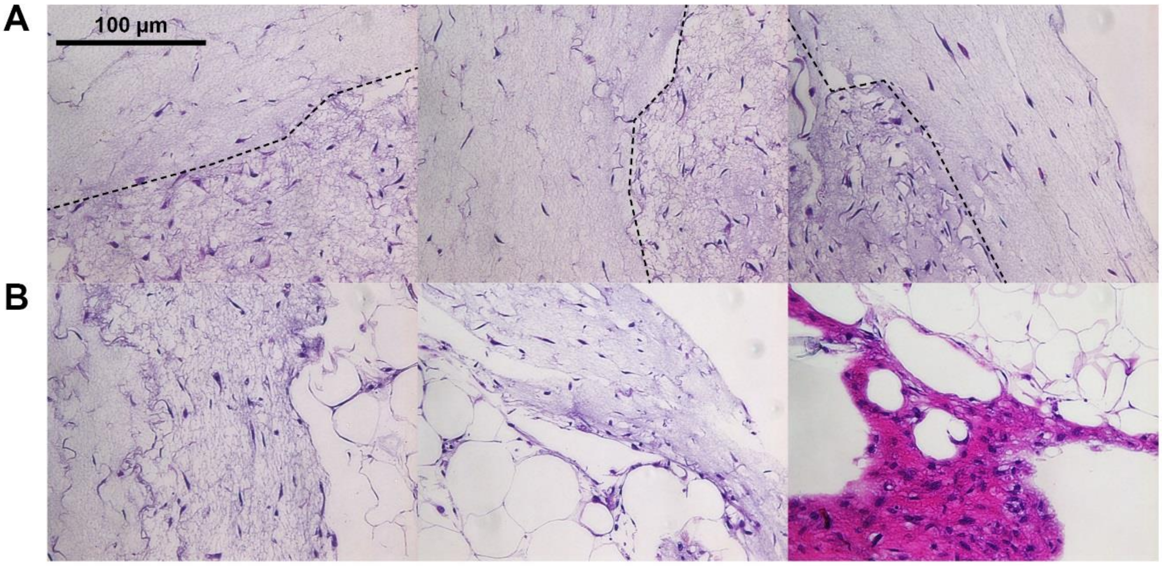


**Supplementary image 2.** Histological sections stained with hematoxylin (blue, stains for nuclei) and eosin (pink, stains for collagen). **(A)** PLMatrix with MSCs that migrated into the hydrogel (the dashed line separates areas where MSCs started to remodel the matrix and areas that are infiltrated by only a few cells). **(B)** Micrographs covering actual areas of MSC outgrowth from adipose tissue into PLMatrix.

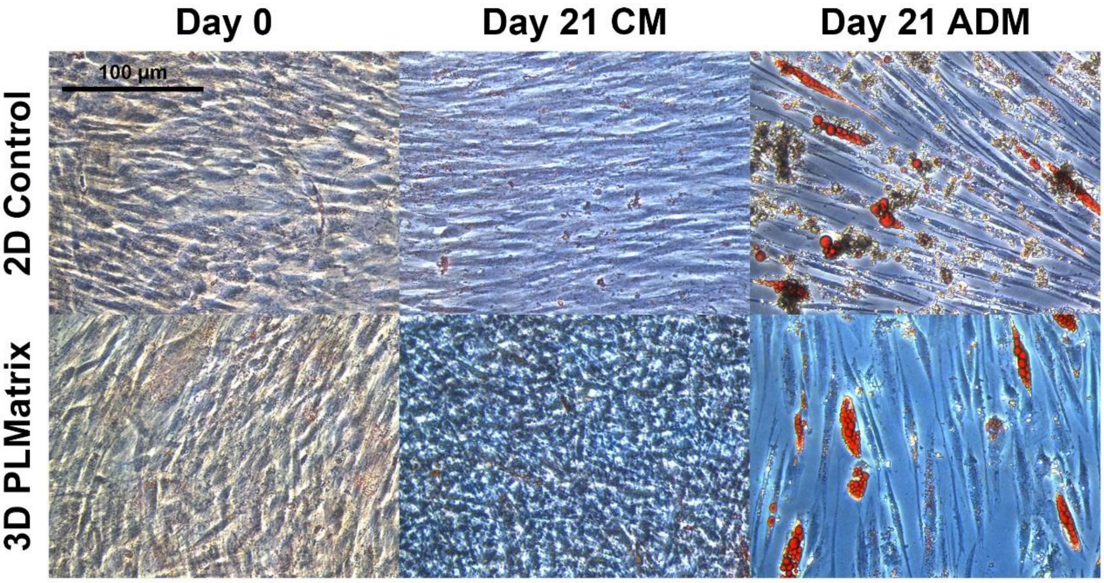


**Supplementary image 3.** Cells derived from isolation by enzymatic treatment (2D control) or from 3D isolation (3D PLMatrix) cultivated for 0 or 21 days in control medium (CM) or adipogenic differentiation medium (ADM) stained with Oil Red O.

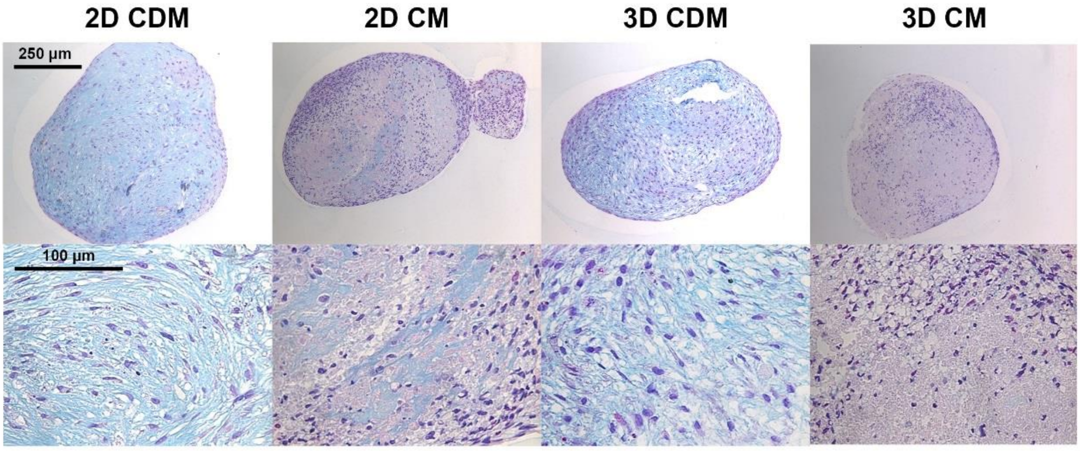


**Supplementary image 4.** Histological sections of cells derived from isolation by enzymatic treatment (2D control) or from 3D isolation (3D PLMatrix) cultivated for 21 days in control medium (CM) or chondrogenic differentiation medium (CDM) stained with Alcian Blue.

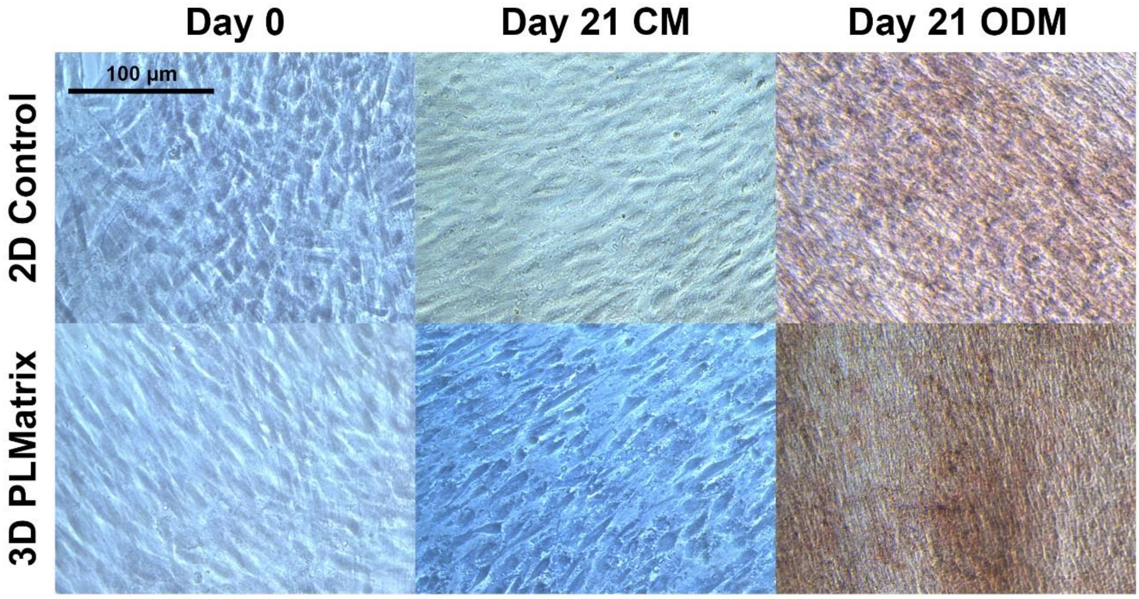


**Supplementary image 5.** Cells derived from isolation by enzymatic treatment (2D control) or from 3D isolation (3D PLMatrix) cultivated for 0 or 21 days in control medium (CM) or osteogenic differentiation medium (ODM) stained with Alizarin Red.

**Supplementary figure 1.** Time series of micrographs of adipose tissue embedded in PLMatrix over a course of 104 h. Cells migrating from the tissue into the matrix can be observed after 72 h.

**Supplementary figure 2.** Histological sections stained with hematoxylin (blue, stains for nuclei) and eosin (pink, stains for collagen). **(A)** PLMatrix with MSCs that migrated into the hydrogel (the dashed line separates areas where MSCs started to remodel the matrix and areas that are infiltrated by only a few cells). **(B)** Micrographs covering actual areas of MSC outgrowth from adipose tissue into PLMatrix.

**Supplementary figure 3.** Cells derived from isolation by enzymatic treatment (2D control) or from 3D isolation (3D PLMatrix) cultivated for 0 or 21 days in control medium (CM) or adipogenic differentiation medium (ADM) stained with Oil Red O.

**Supplementary figure 4.** Histological sections of cells derived from isolation by enzymatic treatment (2D control) or from 3D isolation (3D PLMatrix) cultivated for 21 days in control medium (CM) or chondrogenic differentiation medium (CDM) stained with Alcian Blue.

**Supplementary figure 5.** Cells derived from isolation by enzymatic treatment (2D control) or from 3D isolation (3D PLMatrix) cultivated for 0 or 21 days in control medium (CM) or osteogenic differentiation medium (ODM) stained with Alicarin Red.

**Supplementary video 1.** Cells migrating from adipose tissue into PLMatrix 48 – 104 h after imbedding adipose tissue into PLMatrix.
